# New insights of glycosylation role on variable domain of antibody structures

**DOI:** 10.1101/2021.04.11.439351

**Authors:** Marcella Nunes Melo-Braga, Milene Barbosa Carvalho, Manuela Cristina Emiliano Ferreira, Liza Figueiredo Felicori

## Abstract

N-glycosylation at antibody variable domain (Fv^N-glyco^) has emerged as an important modification for antibody function such as stability and antigen recognition, but it is also associated with autoimmune disease and IgE-mediated hypersensitivity reaction. However, the information related to its role and regulation is still scarce. Therefore, we investigated new insights in this regarding using solved antibodies structures presenting in the Protein Data Bank (PDB). From 130 Fv^N-glyco^ structures, we observed significant findings as a higher prevalence of N-glycosylation in human and mouse antibodies containing IGHV1-8 and IGHV2-2 germline genes, respectively. We also speculate the influence of activation-induced cytidine deaminase (AID) in introducing N-glycosylation sites during somatic hypermutation, specifically on threonine to asparagine substitution. Moreover, we highlight the enrichment of anti-HIV antibodies containing N-glycosylation at antibody variable domain and where we showed a possible important role of N-glycosylation, besides to antigen-antibody interactions, in antibody chain pair and antibody-antibody interactions. These could be a positive secondary effect of glycosylation to enhance antigen binding and further neutralization, including an additional mechanism to form Fab-dimers. Overall, our findings extend the knowledge on the characteristics and diverse role of N-glycosylation at antibody variable domain.

**Key Points:**

- Prevalence of Fv^N-glyco^ in human IGHV1-8 and mouse IGHV2-2 germline genes.
- Enrichment of antibody Fv^N-glyco^ against virus, especially anti-HIV-1.
- Fv^N-glyco^ in the interface with antigen, antibody pair chain, and another antibody.

## Introduction

Antibodies are essential components of the adaptive immune system. They can bind to a great diversity of antigens, and extensive investigations are done to better comprehend their function and to manipulate their properties, enhancing their specificity and affinity for therapeutic purposes (1,2).

To understand the binding mechanism toward antigen neutralization or antibody-induced signaling, a relevant information relies on the study of post-translational modifications (PTMs) that can be intrinsically expressed in antibodies or introduced through engineering. Pyroglutamic acid, disulfide bond, glycation, sulfonation and glycosylation are examples of PTMs present in these humoral components (3,4). Among them, glycosylation is one of the most frequent and complex PTMs that plays an important role in antibody functions and properties (5).

Although there are different types of protein glycosylation, only N-linked and O-linked glycosylation have been reported on antibodies. Glycosylation in the constant Fc region is well characterized in all human antibody isotypes (5,6). The N-glycosylation in this region modulates the Fc-mediated immune response through Fc receptors and complement system protein interaction (3,5). Antibody Fc-glycosylation pattern seems to be tuned during vaccination, for instance, by increasing the level of galactose and sialic acid (7–10), although its exact role in vaccine efficacy is still lacking. Moreover, IgG N297 glycosylation site has been monitored for the onset and progression of autoimmune disease (6,11) and inflammation (12).

Even less known than Fc-glycosylation function is the N-glycosylation characteristics of antibody variable domain (Fv^N-glyco^). Most N-glycosylation encoding sites in this region are introduced during somatic hypermutation (SHM) (13) with few germline genes containing putative sites. Thus, this modification is now considered an additional mechanism to generate antibody diversity together with V(D)J gene rearrangement and somatic hypermutation (14). Interesting to note that 15-25% of human serum IgG may contain Fv^N-glyco^ (15), with higher glycosylation prevalence toward human IgG4 among other subclasses (14). This reinforces their important role for antibody function. Therefore, Fv^N-glyco^ has been associated with: i) modulation of antigen biding (14,16), ii) antibody stability (17), iii) physiological alterations, such as during pregnancy (18), iv) pathophysiological conditions such as immune disease (19,20) and tumor (21), iv) anti-inflammatory effects (22), and also with v) IgE-mediated hypersensitivity reactions (23).

The Protein Data Bank (PDB) (24) is a great source to study the possible role of PTMs as glycosylation in proteins through crystallized structures, including antibodies in complex (or not) with antigens. To retrieve glycan structural information from such structures is a challenge, but the improvement of bioinformatics tools assists in this task (25–27). While there are several studies on antibody Fc glycosylation from PDB structures (28–30), there is no study regarding the Fv^N-glyco^ to the best of our knowledge. In addition, information about Fv^N-glyco^ function and regulation is still scarce. Therefore, we decided to explore PDB available structures using bioinformatic tools to shed light on the antibody Fv^N-glyco^. In this work, we showed a higher prevalence of Fv^N-glyco^ in human IGHV1-8 and mice IGHV2-2 germline genes that contains the N-glycosylation encoded site, although we also speculate a possible role of activation-induced cytidine deaminase (AID) in introducing N-glycosylation sites during SHM. Moreover, we highlight the enrichment of anti-HIV antibodies containing Fv^N-glyco^ and containing different interactions in the analyzed dataset, where we showed a possible important role of N-glycosylation in antibody chain pair and antibody-antibody interactions. These may represent a positive secondary effect of glycosylation that can impact antigen binding and contribute to antibody understanding and engineering.

## Materials and Methods

### Selection of antibodies structures containing glycosylation

To get all Protein Data Bank (PDB) entries that contain at least one antibody variable domain, we inspected the Yvis platform (31) on 24^th^ of July 2020. As a result, we got 4,104 PDB entries with numbered antibody chains, according to the IMGT numbering (32), compared to the entire PDB on the same date (166,891 sequences). Thus, we downloaded each structure file from the Worldwide PDB (wwPDB) server (33). Then, we searched for NAG (N-acetyl-beta-D-glucosamine) in Asn (asparagine) amino acid residue or NAG, MAN (alpha-D-mannose) or BMA (beta-D-mannose) in Ser (serine) or Thr (threonine) amino acid residues on the LINK record of each structure file to investigate N-glycosylation and O-glycosylation, respectively.

Since extracting glycan information from a crystal structure determination process is not a trivial process, we also obtained all the previously antibody structures from the PDB-REDO Databank (34) with the aim to get refined glycan structures (21). Then, we performed the same search based on the LINK record to investigate N- and O-glycosylation of wwPDB-REDO files.

Based on the Yvis platform data and the downloaded structure files, an *in-house* Python script was developed to automatize the process to identify PDB structures with at least one antibody variable domain containing glycosylation and all the analyzed information.

### Extracting information from glycosylated antibodies

The developed script retrieved, for each glycosylation, the molecule type (antibody or antigen), the experimental structure determination method, the resolution, and the glycosylation region sequence (six amino acids before and after the modification). For glycosylated antibodies, the script also informs the modified antibody domain (variable or constant) and for those that contain glycosylation on the variable domain, the script provides: i) the IMGT annotated V and J germline genes, ii) the antibody and protein antigen-producing organism (if the structure presents an antibody in complex), iii) the antigen molecule description, iv) the IMGT position of glycosylation and its region (framework or CDR), and v) the gapped and ungapped variable domain sequence.

We manually inspected the antigen producing organism and molecule description of glycosylated antibodies, comparing it to the literature stated in the corresponding PDB entry. Moreover, we automatically corrected assigned errors as assigned germline genes of chimeric structures and also errors concerning wrong antigen identification, where the carbohydrate was assigned as antigen and not as antibody modification. From PDB structures stated in the literature, we also retrieved the antibody expression system information. From literature and/or from IMGT/3Dstructure-DB (35), we extracted the antibody fragment type (scFv, Fab, or VHH) and, when available, the antibody isotype class and subclass. Finally, because of the variety of antigens found in the PDB entries, we manually grouped them based on the molecule function available on UniProt Knowledgebase (36).

To further evaluate antibody germline preference, modified position and antigen information, we created a non-redundant (NR) set of antibodies by clustering antibody structures that have identical modified variable domain amino acid sequences. The representative structure from each cluster was chosen based on the presence of an antigen, when possible, and better resolution.

### Obtaining glycan structure information from glycosylated antibodies

Due to the difficulty of extracting glycan structural information from such structure files, we submitted all wwPDB or PDB-REDO files containing glycosylation to pdb2linucs, which display the carbohydrate structure in a standardized manner (26,37). In pdb2linucs tool, we selected the options to find carbohydrates in ATOM and HETATM records and, whenever necessary, allowing assigned connections by the distance of the atoms indicated by these same records. We also chose to pdb2linucs to show the carbohydrate structures using the LINUCS (37) notation, favoring the comparison of all analyzed structures. These searches were also implemented in our *in-house* Python script.

### Investigation of glycan interaction partners

To analyze interacting glycans, we extracted all atoms within a maximum 5Å of distance from each asparagine and carbohydrate atoms of Fv^N-glyco^ chains, except for hydrogen. For this, we used the NeighborSearch (38) module of the Biopython toolkit (39) on the PDB-Redo file or wwPDB for those not included in PDB-Redo databank from the entire Fv^N-glyco^ dataset. Then, we defined interaction pairs composed of a glycosylated Asn or its attached carbohydrate of an antibody and another amino acid residue or other molecule of another antibody chain or an antigen. To represent the interactions pairs, we further excluded glycosylated chains from same or different PDBs that contains identical amino acid sequences, and we kept those that presented more interactions.

## Results

### Three percent of analyzed antibody structures present N-glycosylation at the antibody variable domain

Using PDB and PDB-Redo databases as well as an *in-house* computational strategy, we identified a total of 148 PDB entries (231 chains) with glycosylated antibodies (**Supplementary Table 1A**) from 4,104 entries that contain antibody variable domains in complexed or free.

Although we have observed glycosylation in the constant domain of antibody chains (Fab or Fc), most modifications are N-glycosylation in the variable domain (Fv^N-glyco^) (198 modified chains of 131 PDB entries) (Table 1). We also observed only two structures containing O-glycosylation. Interesting to note that one PDB entry (2FBJ) contains N-glycosylation in the variable and constant domain from the same heavy chain.

**Table I.** Overview of PDB entries and antibody chains with glycosylation. The number of occurrences of modified entries/chains are classified by modified residues.

|  | Variable | Constant<sup>#</sup> | Total |
| --- | --- | --- | --- |
| **PDB entries (Modified chains)** | 133 (202) | 16 (29) | 148* (231) |
| **Residue N** | 131 (198) | 16 (29) | 146* (227) |
| **Residue T** | 2 (4) | - | 2 (4) |

<sup>#</sup>Fab or Fc region
\*2FBJ has glycosylation in variable and constant in the same chain

For further analysis, we considered 130 structures with 195 Fv^N-glyco^ chains (**Supplementary Table 1**), after excluding one PDB structure with low resolution (17 Å). The structures in this dataset were solved by X-ray crystallography (96.2%) or cryogenic electron microscopy (cryo-EM) (3.8%) with resolution from 1.6 Å to 6.2 Å (**Supplementary Figure 1A and B**, respectively). However, 80% of the structures have a resolution better than 3.05 Å and the worst resolutions belongs to cryo-EM. Moreover, this set comprises 75 (57.7%) human, 37 (28.5%) mice, 15 (11.5%) chimeric, 2 (1.5%) synthetic and 1 European rabbit (0.8%) antibody structures (**Supplementary Figure 1C**). Although the majority of structures are fragment antigen-binding (Fab) or F(ab′)_2_ (91.5%), there are structures of single-chain variable fragments (scFv) (1.5%) but also structures containing more than one antibody fragment type Fab/scFv (4.6%) or Fab/camelid single-domain antibodies (VHH) (2.3%) (**Supplementary Figure 1D**). Finally, the Fv^N-glyco^ antibodies belong to different isotypes and subclasses: IgG1 (76.9%), IgG2a (5.4%), IgG2 (3.8%), IgG4 (2.3%), IgG and IgA (1.5% each) (**Supplementary Figure 1E**). However, in 8.5% of the structures, no isotype information was retrieved.

Approximately 70% (89 structures) of these structures contain a single modification, but 18%, 7.7% and 5.3% contains 2, 3 and 4 modifications per entry, respectively (**Supplementary Figure 1F**). The reason behind multiple modifications in one structure is a redundancy of variable domain sequence per structure. Because of this, we created a non-redundant (NR) set of antibodies by clustering modified chains that have identical variable domain sequences and choosing one representative sequence per cluster (**Supplementary Table 1A**).

From 195 modified chains, we ended up with a dataset containing 44 clusters (unique modified variable domain sequences) for the heavy chain and 32 clusters for light chains (12 and 20 sequences from lambda and kappa, respectively) (**Table 2**). Only two PDBs entries (4YDJ and 4RNR) have glycosylation in different chains (heavy and light lambda) and no PDB structure has multiple modified sites in the same chain. This non-redundant (NR) set was used to evaluate the germline gene preference, modified position and antigen information.

**Table II.** Summary of PDBs and chains with N-glycosylation at antibody variable domain.

|  | Heavy | Light (kappa) | Light (lambda) | Total |
| --- | --- | --- | --- | --- |
| **Modified chains** | 119 | 49 | 27 | 195 |
| **Modified chains (NR)** | 44 | 20 | 12 | 76 |
| **PDBs entries** | 83 | 32 | 17 | 130* |
| **NR PDB entries** | 43 | 20 | 12 | 73* |

\*4YDJ and 4RNR have N-glycosylation in heavy and light chains
NR: non-redundant set of antibodies

### Predominance of N-glycosylation at human IGHV1-8 and mouse IGHV2-2 germlines

Several IMGT antibodies germlines genes contains N-glycosylation putative site as can be observed by browsing the IMGT/GENE-DB (41). The functional IGHV genes from human antibodies containing N-glycosylation encoded sites are: IGHV1-8 (alleles 1 to 3), IGHV3-13 (alleles 2), IGHV3-33 (allele 2), IGHV4-34 (alleles 1 to 11 and 13), IGHV5-10-1 (alleles 1 to 4), IGKV5-2 (allele 1), IGLV3-22 (alleles 1 and 3) and IGLV5-37 (allele 1). While the functional genes from mice antibodies are IGHV1-22 (allele 1), IGHV1-49 (allele1), IGHV1S29 (alleles 1 and 2), IGHV2-2 (alleles 2 and 3), IGHV2-6-3 (allele1), IGHV4-1 (allele 2), IGHV4-2 (allele 2), IGKV20-101-2 (allele 1), IGK5-48 (allele 1) and IGKV8-24 (allele 1).

Even with a small NR set, we decided to investigate the predominance of germline genes of Fv^N-glyco^ sequences for their variable (V) and joining (J) genes, including their alleles. From 76 NR chains, 49 were assigned to human germline genes, 26 to mouse and one to rabbit. Since only one N-glycosylated chain from rabbit was found, this germline gene was not represented in Figure 1. The 49 human sequences belong to 15 different IGHV (18 alleles), 9 IGKV and 8 IGLV and 6 IGHJ, 5 IGKJ and 4 IGLJ germline genes (**Figure 1A and 1B**). The most represented germline genes containing N-glycosylation sites in this dataset are the IGHV1-8, IGHV3-23, IGLV2-14, IGKV1-33 and IGKV3-11. Interesting to note, that from 18 IGHV different alleles containing N-glycosylation, only 3 of them have the encoded N-X-S/T putative site. Regarding the J genes, the most frequents are IGHJ4, IGLJ2 and IGKJ1, together with IGKJ5.

**Figure 1.**
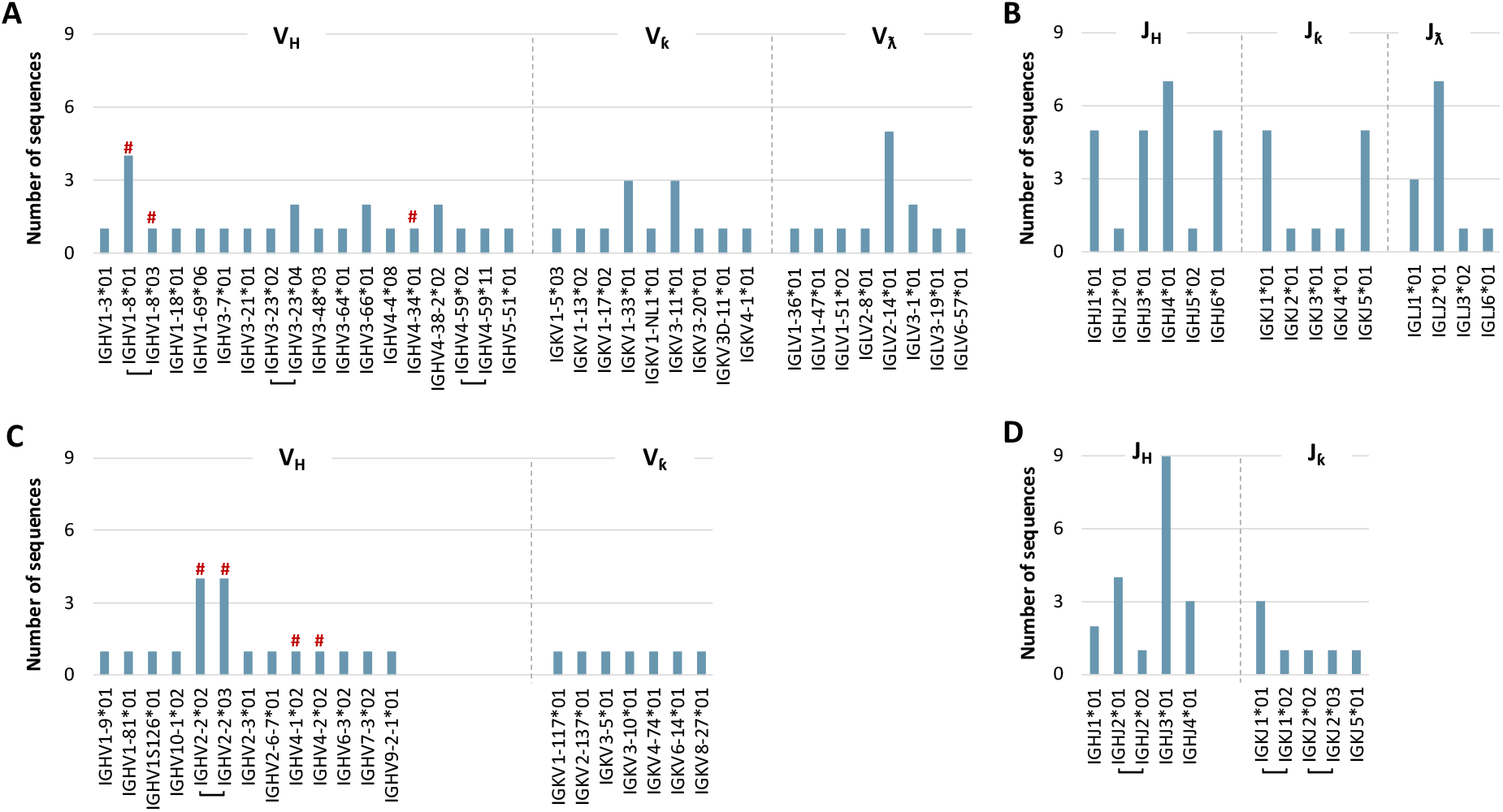
Germline of the N-glycosylated antibodies variable domain. Human germline alleles of (A) variable and (B) joining genes from heavy, kappa and lambda chains. Mouse germline alleles of (C) variable and (D) joining genes from heavy and kappa chains. ^#^N-glycosylation germline-encoded alleles. Square brackets indicate different alleles from the same gene.

Also, the 26 mouse sequences were distributed within 12 IGHV (13 alleles) and 7 IGKV germline genes, as well as 4 IGHJ and 3 IGKJ genes (**Figure 1C and 1D**). There is only one abundant variable germline gene (IGHV2-2), and the more abundant joining genes from heavy and kappa chains are IGHJ3 and IGKJ1 (including two alleles), respectively.

Considering the amino acid mutation in the N-glycosylation motif (NXS/T) from 76 NR sequences, we observed only 14 sequences with germline genes encoded sites belonging to human IGHV1-8 and mouse IGHV2-2, IGHV4-1*02 and IGHV4-2*02 genes/alleles. It was also observed changes in all three amino acids positions from the motif, with the first position containing the highest variation when comparing the dataset sequences with their corresponding germline sequences (**Figure 2A**). This data reinforces the importance of SHM in introducing Fv^N-glyco^. Interestingly, from 69 sequences (91%) containing a mutation at V genes (**Figure 2B**), 50 of them alters the first amino acid with a higher substitution from polar (specially Thr and Ser) to a Asn (**Figure 2C**). Thus, this result implies the maintenance of hydrophilic characteristics with glycosylation.

**Figure 2.**
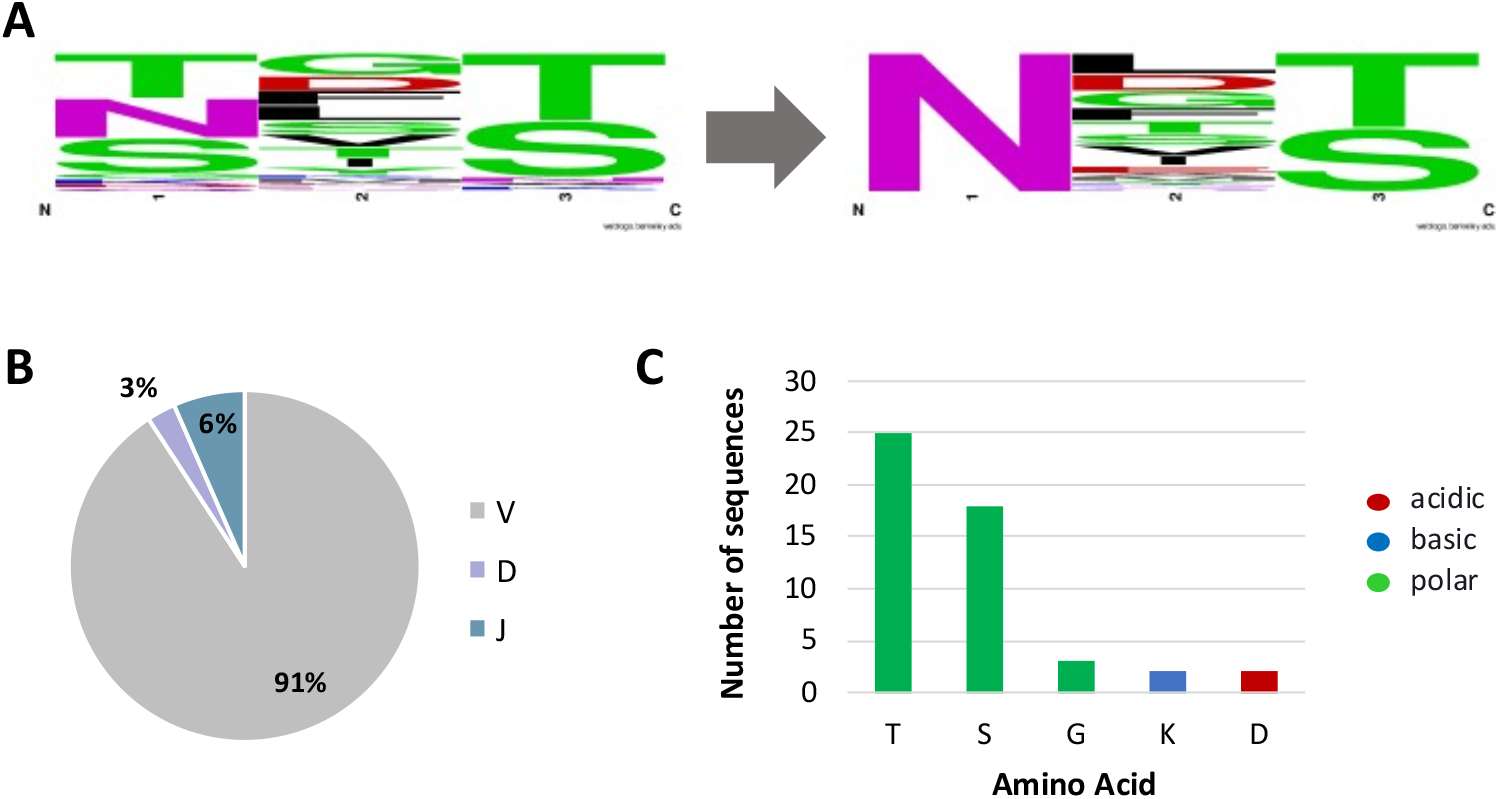
Mutations in the germline N-glycosylation motif. (A) WebLogo (https://weblogo.berkeley.edu/) representation of alteration in the N-glycosylation motif in the germline and those modified sequences from the NR dataset containing 76 sequences. (B) Distribution of glycosylation across germline variable V, diversity D and joining J genes. (C) Amino acids present in the germline V gene sequences preferred substituted in the first position of N-glycosylation motif by asparagine - N.

### Heterogeneity of glycosylation structures found mostly on FR1 and FR3

We next extended our analysis to identify the glycosylation preference region, considering our NR set. We observed that FR3 (including DE loop) contains the majority of Fv^N-glyco^ in heavy chain (23 sequences) and kappa chain (9 sequences) followed by FR1 in both light kappa and lambda chains (7 and 10 seq, respectively). Also, we observed glycosylation in all loop regions (CDRs and DE) in the heavy chain, while only CDR1 and DE loops in kappa and CDR3 in lambda chains are glycosylated (**Figure 3**).

**Figure 3.**
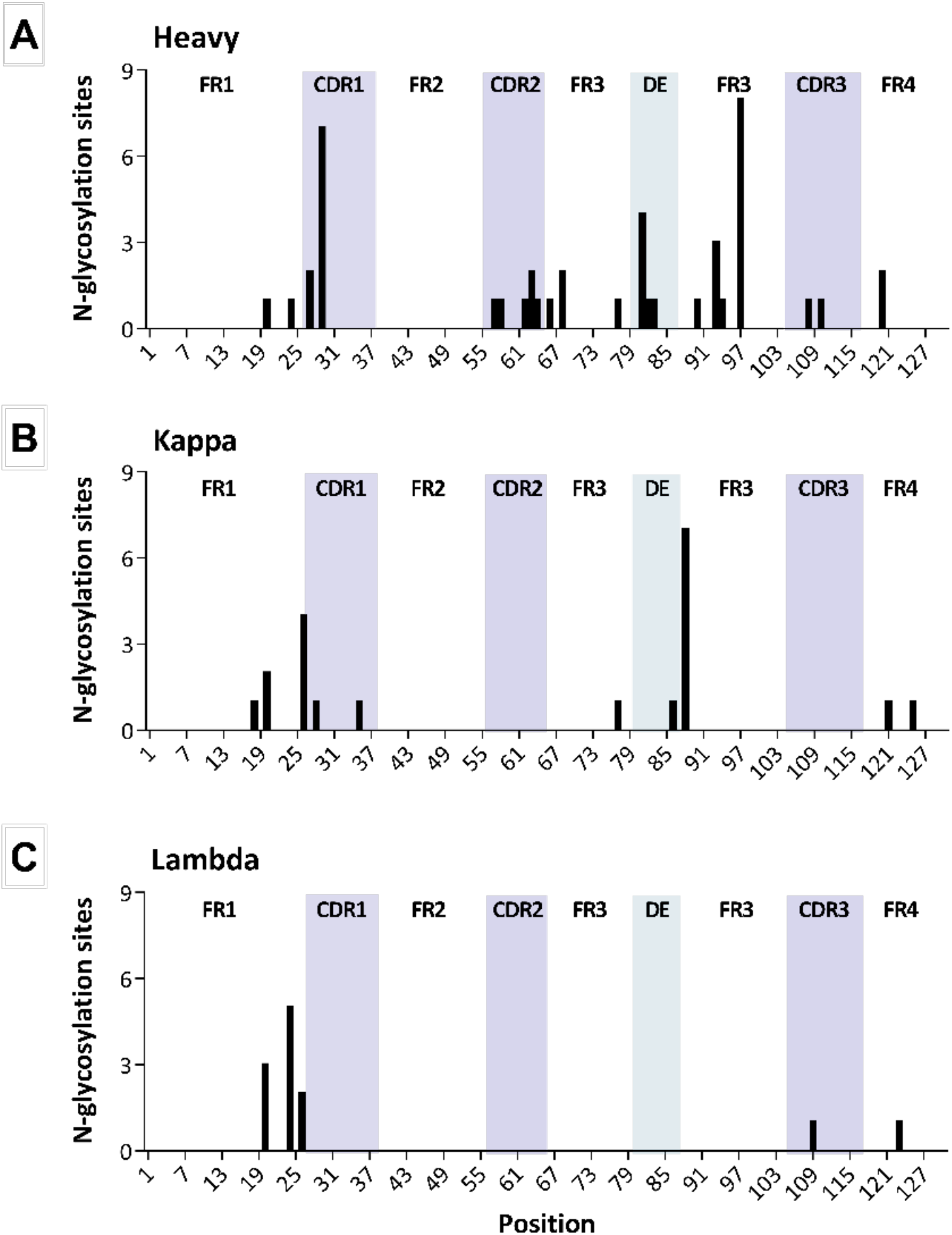
Number of N-glycosylation sites distributed in the antibody variable region from the non-redundant dataset. IMGT position of N-glycosylation in the (A) heavy chain, and light (B) kappa and (C) lambda chains. Loop regions are highlighted.

To further evaluate the glycan structures on the antibody variable domain, we considered all 195 modified chains since same sequences could have different attached glycans. Therefore, we mapped the glycan structures presented in each variable domain position, where we observed a prevalence of only one or two N-acetylglucosamine (GlcNAc also known as NAG) with fewer paucimannose and complex biantennary glycan structures containing up to 10 carbohydrates (**Figure 4**). Besides, glycan structures with fucose (FUC) are highly abundant, whereas only two was identified with sialic acid (SA). This result can partially be explained by different host antibody expression and glycosidase treatment of antibodies before crystallization.

**Figure 4.**
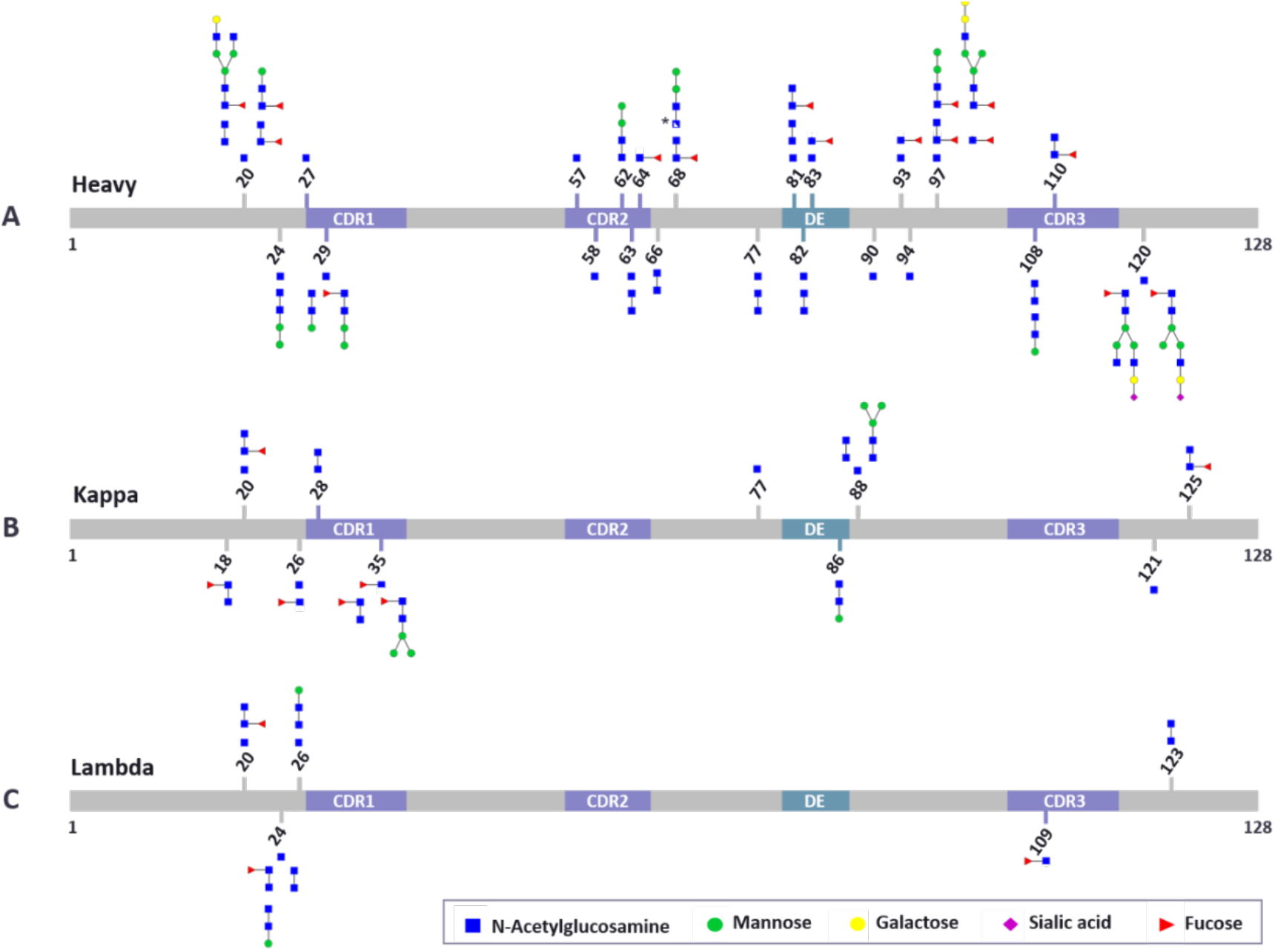
Glycan structures in the antibody variable domain. Heterogeneity of non-redundant glycosylation structure per IMGT position distributed across the variable domain of heavy (A), kappa (B) and lambda (C) chains. Some positions may contain different glycan structures when different chains present glycosylation in same position (e.g., heavy 20, kappa 35, lambda 24, among others).

Interesting, the heavy chain position 97 (FR3) (**Figure 3A**) contains the higher number of N-glycosylated sequences and also higher glycan heterogeneity (**Figure 4**) besides all of them come from mouse IGHV2-2 germline gene that contains the encoded N-glycosylation site.

### Anti-HIV variable chain antibodies are enriched in N-glycosylation

Next, we investigated whether antibody Fv^N-glyco^ could be subject to antigen preference, representing a possible selection mechanism in such response. Considering our NR set, most of the glycosylated entries contain antigen-antibody complexes (54 PDB entries representing 73.9%). In this work, we retrieved antigen source for these complexed antibodies and also for the remaining free antibodies (26% - 19 PDB entries) (**Figure 4A**). Although antibodies against virus species are overrepresented in our set, we also observe antigens from parasites, mammalians, bacteria, and molecules such as hapten and carbohydrates. However, it is clear the predominance of antigens from human immunodeficiency virus 1 (HIV) and humans among the antigens from 17 different organisms or non-protein antigens detected in this set (**Figure 5A**). From 73 PDB structures with glycosylated antibody chains, 28 are from antibodies against HIV proteins, representing 38% of structures, while 20 (27%) are against human proteins.

**Figure 5.**
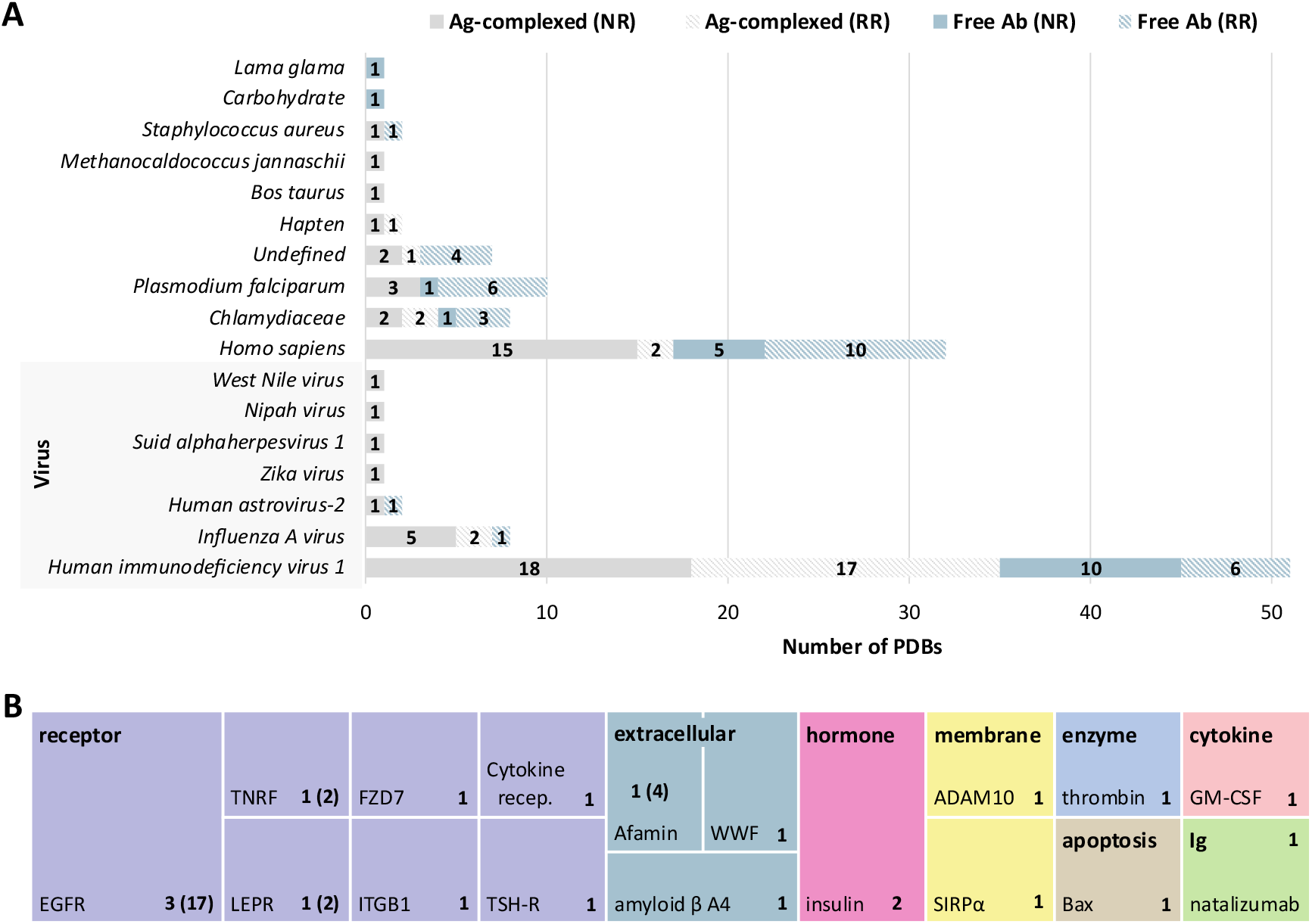
Antigen diversity of PDB entries containing N-glycosylation at antibody variable domain. (A) Number of PDB entries classified by antigen-producing organism or non-protein antigen (ex. hapten and carbohydrate) from non-redundant (NR) dataset (73 PDBs in solid fill) and the removed redundant (RR) in light pattern filled, totalizing 130 PDB entries. (B) Distribution of antigen molecules from different human protein classes, where box areas are proportional to the number of the non-redundant PDBs entries. The box also contains the number of total redundant structures between brackets. Colors identify different antigen molecule classification. EGFR: epidermal growth factor receptor; TNFR: tumor necrosis factor receptor; FZD7: frizzled-7; ITGB1: integrin beta-1; LEPR: leptin receptor; TSH-R: thyrotropin receptor; WWF: von Willebrand factor; ADAM10: disintegrin and metalloproteinase domain-containing protein10; SIPRα: signal regulatory protein α; GM-CSF: Granulocyte-macrophage colony-stimulating factor; Ig: immunoglobulin.

On the other hand, as expected, there is a greater variety of antigen types from human belonging to 20 PDB entries in the NR set. They are receptors (35%), extracellular proteins (15%), membrane proteins (10%), hormones (5%), cytokines (5%), enzymes (5%), and even an antibody (5%) (**Figure 5B**).

Within the whole antibody database where the structures were extracted - Yvis (31) - the complexed antibodies represent 70% of structures, where 348 and 733 structures are in complex with molecules from HIV and humans, respectively. Comparing only antibody structures in complex with HIV molecules from our entire N-glycosylation dataset (35 structures, **Figure 5A**) and the Yvis database, approximately 10% of these antibodies contain Fv^N-glyco^. Remarkably, this comparison pointed to a possible significant impact of antibody glycosylation against other virus organisms such as Human astrovirus-2 (100%), Suid alpha-herpesvirus 1 and Nipah virus (50% each) and West Nile virus (16.7%). These results imply a positive binding effect of Fv^N-glyco^ and antigens from virus.

Regarding human antigens, the growth factor receptor (EGFR) was the most targeted in our dataset. This receptor is targeted by the well-known therapeutic antibody Cetuximab, a glycosylated chimeric monoclonal antibody used to treat colorectal cancer patients (42). Although there are 18 antibody-EGFR complex structures in the Yvis database, with 13 of them presenting N-glycosylation putative site, only four contain N-glycosylation at the variable domain (16.7%), even though the Cetuximab is known to be glycosylated at position 88 or 97 according to IMGT numbering. This example suggested a mutation in the other structures and a possible deglycosylation before crystallization.

### N-glycosylated antibodies site interacts preferentially with an antibody chain or a different antibody

Next, we investigated the possible interactions of the antibody variable domain modified site and their glycan structures by extracting all residues within 5Å of distance from them. Since heterogeneous glycan structures could lead to different interactions, we perform this analysis with all 130 PDB entries (195 modified chains) from our entire N-glycosylation dataset. Thus, we identified 7, 10 and 8 entries with antibody-antigen (Ab-Ag), antibody-chains pair (Ab-chain pair) and antibody-antibody (Ab-Ab) interactions, respectively (**Supplementary Table 1_B**). It is interesting to note that antibodies against peptides/proteins of HIV and *Plasmodium falciparum* contain all three different interaction types. **Figure 6** shows an example of each interaction on anti-HIV antibodies. In addition, the same structure could have more than one interaction, such as 6UUD that had both Ab-Ag and Ab-chains pair interactions while 2G5B had Ab-chain pair and Ab-Ab interactions. There are also examples of different Ab-Ab interactions, as we observed in the 3MUG entry that contains interaction between light chains and light-heavy chains.

**Figure 6.**
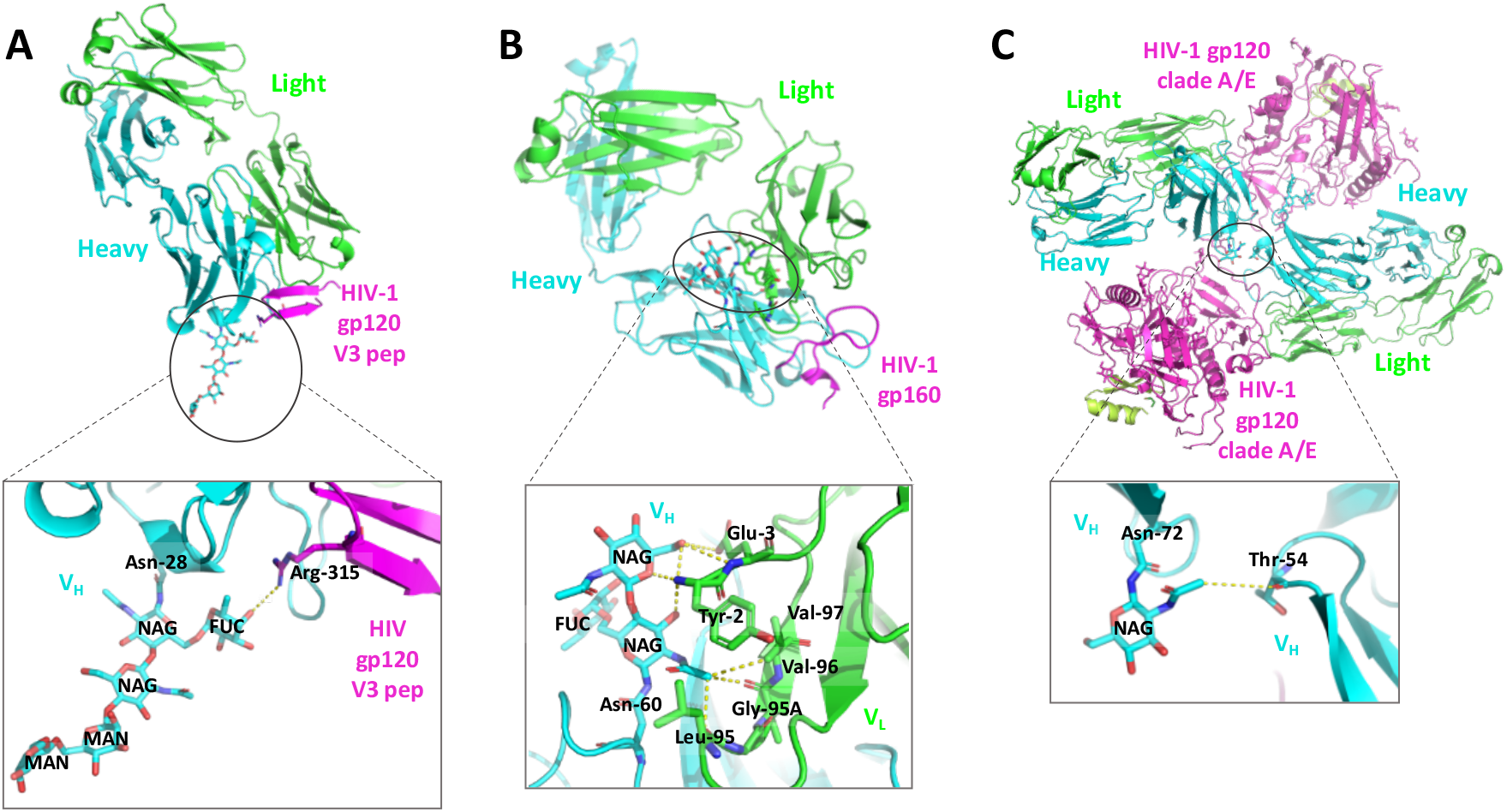
Antibody N-glycosylation in interaction with antigen, antibody chain pair and another antibody. Ribbon representation of PDB structures (with antibody light chain in green, antibody heavy chain in blue and antigen in pink) together with the zoom view of the interaction interface containing the glycosylated antibody variable domain site. (A) Interaction between fucose (FUC) from glycan structure of heavy chain with Arg-315 from HIV antigen in PDB 6DB6. (B) Interaction of two NAGs from heavy chain with Tyr-2, Glu-3, Leu-95, Gly-95A, Val-96 and Val-97 of the light chain pair in PDB 3UJI. (C) Interaction of NAG from antibody heavy chain with Thr-54 from other antibody heavy chain in PDB 6MFP. All examples are antibodies complexed with HIV-1 molecule. Figures were created in PyMOL software.

**Figure 7** shows the number of NR interactions from antibody variable domain modified asparagine and the carbohydrates attached to its structure with residues from the antigen (Ab-Ag), light/heavy chain (Ab-chain pair) and another antibody (Ab-Ab). We observed that modified asparagine interacts more with alanine, aspartic acid and tyrosine from Ab-chain pairs (**Figure 7**). Moreover, NAG interacts with a wide variety of residues, showing higher interaction among polar residues serine (7) and glycine (4) as well as charged residues in Ab-Ab. Considering the residues from Ab-chain pair, NAG interacts more with polar residues tyrosine (5) and glycine (3), acidic residue glutamic acid (3), neutral residue asparagine (3), and hydrophobic residue proline (3). Also, NAG interacts more with basic residue lysine (3) from Ag molecules. MAN interacts mainly with residues from other Ab, especially polar threonine (2), but we also observed interactions with residues from ag and ab-chain pair. On the other hand, FUC interacts more with neutral glutamine (2) from Ab-chain pair residues and polar glycine (2) from Ab-Ab, but also interacts with residues from Ag molecules (**Figure 7**). In general, water mediated interaction was also observed (**Supplementary Table 1_B**).

**Figure 7.**
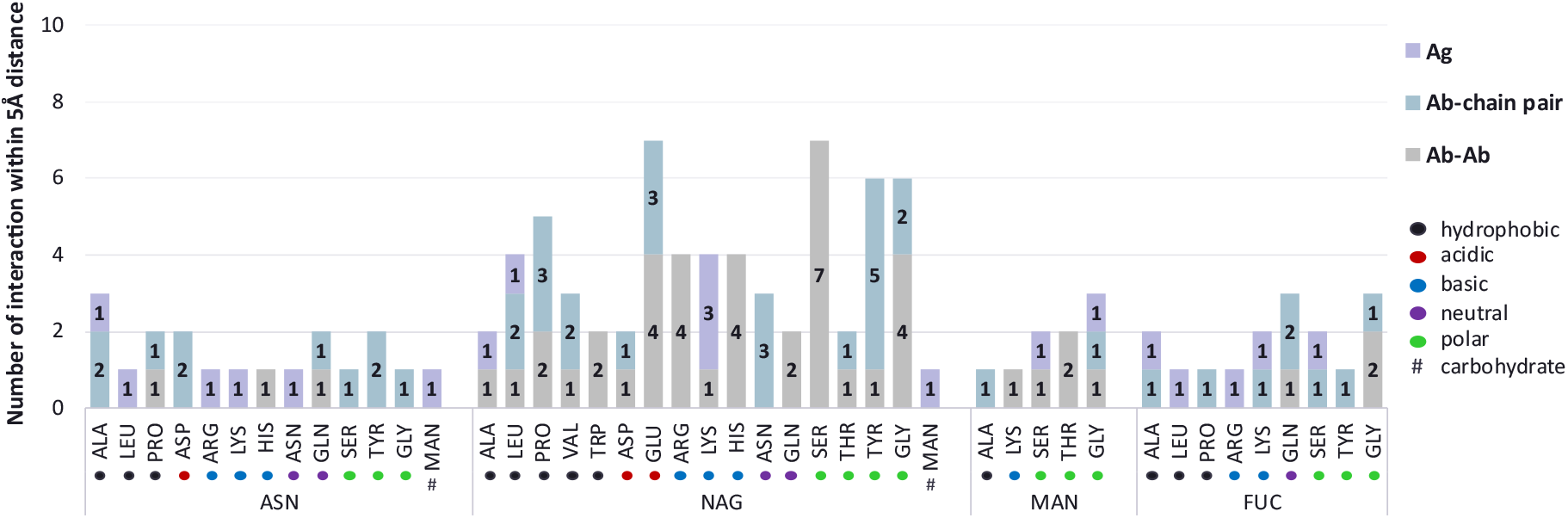
Interactions of antibody variable domain modified sites and its N-glycans within 5Å distance. Number of interactions of glycosylated asparagine (Asn) and its attached glycan with residues from antigen (Ag), antibody (Ab) chain pair or another Ab (Ab-Ab). Color balls represent the interacting residues according to their amino acid classification or # for carbohydrates. ASN (asparagine), NAG (N-acetylglucosamine), MAN (Mannose), FUC (fucose).

## Discussion

In the present study, we characterized Fv^N-glyco^ using solved antibodies structures. We found that 3% of antibody structures contain glycosylation at variable domain, a similar percentage of general glycosylated proteins found in the entire PDB databank (43). We believe this number is underestimated since the crystallization of glycoproteins is challenging, and a deglycosylation step is often performed before it to improve crystal quality (44,45). Thus, a partial deglycosylation may explain the short glycans found in most of the analyzed structures. In addition, the use of a bacterial antibody expression system can be responsible for this underrepresented number of glycosylated antibodies in the Protein Data Bank. The host expression can also impact the glycan repertoire contributing to the complexity of this modification (46). Besides that, some of the human cell lines used for antibody expression lack N-acetylglucosaminyltransferase I (GnTI) activity and therefore lack complex N-glycans, also observed in this study. Another reason would be the preference in studying IgG1 antibodies. Although all human IgG subclasses contain glycosylation on the variable domain, IgG4 has a higher prevalence (10), represented by only 2.3% of our dataset structures. Also, we suspect that this underestimation can be due to the difficulty in interpreting the carbohydrate on X-ray diffraction data and glycan heterogeneity.

Additionally, we observed higher glycosylation frequencies for mouse IGHV2-2 and human IGHV1-8 that contains putative N-glycosylated encoded sites. Although the percentage of structures containing IGHV1-8 was lower in the Yvis antibody database (0.4%) and also relatively lower within human antibody repertoire studies ranging from 1.1-5% (47,48), this germline seems to have a higher glycosylation occupancy representing 70% of those sequences containing N-glycosylation motif. Important to mention that we also found one sequence that lost the germline IGHV1-8 encoded motif. Conversely to IGHV1-8, the germline IGHV4-34 is commonly found without glycosylation at the putative site (49). Accordingly, we observed in our dataset a glycosylation in a different position of the encoded site.

Several identified germline genes have no encoded putative N-glycosylation site, but previous studies showed that the mutation of a single nucleotide during SHM could create an N-glycosylation motif, suggesting a preset position to quickly introduce N-glycosylation (14). Indeed, more than 78% of these SHM putative sites was observed in switched memory B cells on healthy human antibody repertoire studies (14). These single nucleotide changes also explain some of our observation, where we found a higher mutation from threonine and serine into an asparagine to introduce the N-glycosylation motif. Although these mutations are considered missense, they maintained the hydrophilic feature considering the glycosylation.

Previous work showed most of the putative N-glycosylation encoded germline sites and those introduced during SHM are preferentially in or near the loop regions, and most of them are occupied (14,50). Although we identified glycosylation in the majority of antibody regions, including all fours loops (CDR1, CDR2, CDR3 and DE) and frameworks (FR1, FR3, FR4), most of glycosylation are in FR3 and FR1. Whether one believe activation-induced cytidine deaminase (AID) enzyme plays a role in increasing N-glycosylation at antibody variable domain, it requires further investigation.

Strikingly, there is no glycosylation placed in the FR2 region, also observed by previous studies with no or fewer putative N-glycosylation in this region (14,50,51). One of the reasons could be the internal position of FR2 in the variable domain (52) or even the lower intrinsic mutation rate in this region on heavy chain. However, in the light chains, this region could have a higher mutation frequency than CDR2 (53). Moreover, a relative deleterious effect on antibody structure, antigen-binding, or antibody secretion was observed with mutagenesis studies in the FR2 region (54).

Another important finding was the higher number of glycosylated antibodies against virus organisms, especially HIV-1, which is the virus with more complexed structures in the Yvis database. Not surprisingly, all HIV antigens found in our study are the envelope proteins, mainly gp120 and its precursors gp160 and gp140, since these molecules are the principal targets for anti-HIV treatment (55). When normalization was performed between the complexed structures from our dataset and the Yvis database, 10% of HIV-complexed antibodies contains Fv^N-glyco^. We believe this number is also underestimated for the same reasons already pointed before. Remarkably, anti-HIV antibodies were also enriched when we evaluate the interaction of the modified site and its glycan structures. Here, we showed that N-glycosylation plays a role in antigen, antibody chain pair and between other antibody interactions. From 23 structures containing one or more types of these interactions, 47.8% are anti-HIV antibodies, contributing to 50% of structures with Ab-chain pair interaction, 50% Ab-Ab interactions, and 28.6% antigen interactions. Several of Ab-Ab interactions occurs between the same chain type from other antibody copy but also between different chain types. Indeed, Fab-dimerization, also referred as domain-swapped dimer, has been shown to enhance HIV neutralization, in glycan-reactive antibodies that recognizes high-mannose glycan on the HIV-1 envelope (56,57). To the best of our knowledge, Fv^N-glyco^ has not been implied as an additional mechanism of Fab-dimerization as our observations may suggest. In this case, the N-glycosylation in Ab-chain pair and Ab-Ab may represent a positive secondary effect on antigen binding, which is also supported by our work since complexed structures contain these types of interactions. However, this Ab-Ab interaction could also be an auto-anti-idiotype, since glycans at loop regions were observed.

In general, we observed that glycosylation at all loop regions (CDR1, CDR2, CDR3 and DE) was involved in Ag interaction, while at FR3, FR4 and CDR3 was involved in Ab-chain pair, and FR1, FR3, FR4 but also loops (CDR1, CDR3 and DE) was involved in Ab-Ab interaction. Surprisingly, the first NAGs, followed by the second NAG from glycan structure was essential for the latest interactions. Although we showed important effects of glycosylation, there are divergent observations among the antigen-binding or neutralization. While Lee *et al* (58) and Pancera *et al* (59) reported a non-disturbance of antigen interaction or neutralization by removing the glycan structure, Romijn *et al* (60) showed the direct interaction of antibody glycosylation with the antigen. However, it is important to mention that in the study of Pancera *et al* (59), a minimal glycan structure (one NAG) was still present and in our believe only a mutation site or complete deglycosylation would rule out the non-effect of glycosylation. Moreover, glycoengineering can enhance antigen-binding affinity in 50-fold by introducing N-glycosylation on antibody variable domain (61).

Despite the fact that we showed new direction on the role of N-glycosylation at antibody variable domain, further investigations are necessary. This includes studies with fully glycosylated structures and completely/partially deglycosylation or site directed mutagenesis to address the raising questions. Moreover, crystallography may not represent the conformation of molecules in a physiological state, especially those containing glycosylation, since it does not consider glycan flexibility. Therefore, crystallography combined with liquid phase assays, as nuclear magnetic resonance (NMR), could improve the comprehension of the role of Fv^N-glyco^. In addition, a combination of immunoglobulin repertoire sequencing (e.g., RNAseq) with glycoproteomics studies could improve the knowledge behind germline genes and position preference of glycosylation in antibodies. Together this information will contribute to antibody engineering to develop antibodies with higher affinity, selectivity, stability, and lower side effects.

## Acknowledgements

We thank Professor Lucas Bleicher for suggesting the use of PDB-REDO and Professor Giuseppe Palmisano for guidance and discussions. We also thank Lucas Silva for designing abstract figure.

## Disclosures

The authors have no financial conflicts of interest

## Footnotes

This study was supported by a PNPD postdoctoral fellowship Coordenação de Aperfeiçoamento de Pessoal de Nível Superior – Brazil (CAPES) and also for the grants [grant numbers 88887.506611/2020-00, 88887.504420/2020-00 and 935/19 (COFECUB)]; Fundação de Amparo a Pesquisa de Minas Gerais (FAPEMIG) [grant numbers PPM-00615-18, Rede Mineira de Imunobiologicos grant #REDE-00140-16]; Conselho Nacional de Desenvolvimento Científico e Tecnológico (CNPq) [Pq to LFF];

## Abbreviations

Ab: antibody
Ag: antigen
AID: Activation-induced cytidine deaminase
Fab: Ag-biding fragment
Fv^N-glyco^: glycosylation at antibody variable domain
NR: no-redundant
V(D)J: variable diversity joining
SHM: somatic hypermutation

## Notes

### Competing Interest Statement

The authors have declared no competing interest.

## References

1. Lu RM, Hwang YC, Liu IJ, Lee CC, Tsai HZ, Li HJ, et al. Development of therapeutic antibodies for the treatment of diseases. Journal of Biomedical Science. 2020.

2. Elgundi Z, Reslan M, Cruz E, Sifniotis V, Kayser V. The state-of-play and future of antibody therapeutics. Advanced Drug Delivery Reviews. 2017.

3. Higel F, Seidl A, Sörgel F, Friess W. N-glycosylation heterogeneity and the influence on structure, function and pharmacokinetics of monoclonal antibodies and Fc fusion proteins. European Journal of Pharmaceutics and Biopharmaceutics. 2016.

4. Liu H, Ponniah G, Zhang HM, Nowak C, Neill A, Gonzalez-Lopez N, et al. In vitro and in vivo modifications of recombinant and human IgG antibodies. mAbs. 2014.

5. Arnold JN, Wormald MR, Sim RB, Rudd PM, Dwek RA. The impact of glycosylation on the biological function and structure of human immunoglobulins. Annual Review of Immunology. 2007.

6. Maverakis E, Kim K, Shimoda M, Gershwin ME, Patel F, Wilken R, et al. Glycans in the immune system and The Altered Glycan Theory of Autoimmunity: A critical review. Journal of Autoimmunity. 2015.

7. Wang TT, Maamary J, Tan GS, Bournazos S, Davis CW, Krammer F, et al. Anti-HA Glycoforms Drive B Cell Affinity Selection and Determine Influenza Vaccine Efficacy. Cell. 2015;

8. Mahan AE, Jennewein MF, Suscovich T, Dionne K, Tedesco J, Chung AW, et al. Antigen-Specific Antibody Glycosylation Is Regulated via Vaccination. PLoS Pathog. 2016;

9. Selman MHJ, De Jong SE, Soonawala D, Kroon FP, Adegnika AA, Deelder AM, et al. Changes in antigen-specific IgG1 Fc N-glycosylation upon influenza and tetanus vaccination. Mol Cell Proteomics. 2012;

10. Vestrheim AC, Moen A, Egge-Jacobsen W, Reubsaet L, Halvorsen TG, Bratlie DB, et al. A pilot study showing differences in glycosylation patterns of igg subclasses induced by pneumococcal, meningococcal, and two types of influenza vaccines. Immun Inflamm Dis. 2014;

11. Šimurina M, de Haan N, Vučković F, Kennedy NA, Štambuk J, Falck D, et al. Glycosylation of Immunoglobulin G Associates With Clinical Features of Inflammatory Bowel Diseases. Gastroenterology. 2018;

12. Plomp R, Ruhaak LR, Uh HW, Reiding KR, Selman M, Houwing-Duistermaat JJ, et al. Subclass-specific IgG glycosylation is associated with markers of inflammation and metabolic health. Sci Rep. 2017;

13. Dunn-Walters D, Boursier L, Spencer J. Effect of somatic hypermutation on potential N-glycosylation sites in human immunoglobulin heavy chain variable regions. Mol Immunol. 2000;

14. Van De Bovenkamp FS, Derksen NIL, Ooijevaar-de Heer P, Van Schie KA, Kruithof S, Berkowska MA, et al. Adaptive antibody diversification through N-linked glycosylation of the immunoglobulin variable region. Proc Natl Acad Sci U S A. 2018;

15. van de Bovenkamp FS, Hafkenscheid L, Rispens T, Rombouts Y. The Emerging Importance of IgG Fab Glycosylation in Immunity. J Immunol. 2016;

16. Leibiger H, Wüstner D, Stigler RD, Marx U. Variable domain-linked oligosaccharides of a human monoclonal IgG: Structure and influence on antigen binding. Biochem J. 1999;

17. van de Bovenkamp FS, Derksen NIL, van Breemen MJ, de Taeye SW, Ooijevaar-de Heer P, Sanders RW, et al. Variable domain N-linked glycans acquired during antigenspecific immune responses can contribute to immunoglobulin G antibody stability. Front Immunol. 2018;

18. Bondt A, Rombouts Y, Selman MHJ, Hensbergen PJ, Reiding KR, Hazes JMW, et al. Immunoglobulin G (IgG) fab glycosylation analysis using a new mass spectrometric high-throughput profiling method reveals pregnancy-associated changes. Mol Cell Proteomics. 2014;

19. Sabouri Z, Schofield P, Horikawa K, Spierings E, Kipling D, Randall KL, et al. Redemption of autoantibodies on anergic B cells by variable-region glycosylation and mutation away from self-reactivity. Proc Natl Acad Sci U S A. 2014;

20. Youings A, Chang SC, Dwek RA, Scragg IG. Site-specific glycosylation of human immunoglobulin G is altered in four rheumatoid arthritis patients. Biochem J. 1996;

21. Coelho V, Krysov S, Ghaemmaghami AM, Emara M, Potter KN, Johnson P, et al. Glycosylation of surface Ig creates a functional bridge between human follicular lymphoma and microenvironmental lectins. Proc Natl Acad Sci U S A. 2010;

22. Käsermann F, Boerema DJ, Rüegsegger M, Hofmann A, Wymann S, Zuercher AW, et al. Analysis and functional consequences of increased fab-sialylation of intravenous immunoglobulin (IVIG) after lectin fractionation. PLoS One. 2012;

23. Chung CH, Mirakhur B, Chan E, Le Q-T, Berlin J, Morse M, et al. Cetuximab-Induced Anaphylaxis and IgE Specific for Galactose-α-1,3-Galactose. N Engl J Med. 2008;

24. Berman HM, Westbrook J, Feng Z, Gilliland G, Bhat TN, Weissig H, et al. The Protein Data Bank. Nucleic Acids Research. 2000.

25. Van Beusekom B, Lütteke T, Joosten RP. Making glycoproteins a little bit sweeter with PDB-REDO. Acta Crystallogr Sect F Struct Biol Commun. 2018;

26. Lütteke T, Frank M, Von Der Lieth CW. Data mining the protein data bank: Automatic detection and assignment of carbohydrate structures. In: Carbohydrate Research. 2004.

27. Joosten RP, Lütteke T. Carbohydrate 3D structure validation. Current Opinion in Structural Biology. 2017.

28. Borrok MJ, Jung ST, Kang TH, Monzingo AF, Georgiou G. Revisiting the role of glycosylation in the structure of human IgG Fc. ACS Chem Biol. 2012;

29. Krapp S, Mimura Y, Jefferis R, Huber R, Sondermann P. Structural analysis of human IgG-Fc glycoforms reveals a correlation between glycosylation and structural integrity. J Mol Biol. 2003;

30. Ahmed AA, Giddens J, Pincetic A, Lomino J V., Ravetch J V., Wang LX, et al. Structural characterization of anti-inflammatory immunoglobulin G Fc proteins. J Mol Biol. 2014;

31. Carvalho MB, Molina F, Felicori LF. Yvis: Antibody high-density alignment visualization and analysis platform with an integrated database. Nucleic Acids Res. 2019;

32. Lefranc MP, Pommié C, Ruiz M, Giudicelli V, Foulquier E, Truong L, et al. IMGT unique numbering for immunoglobulin and T cell receptor variable domains and Ig superfamily V-like domains. Dev Comp Immunol. 2003;

33. Berman H, Henrick K, Nakamura H. Announcing the worldwide Protein Data Bank. Nature Structural Biology. 2003.

34. Joosten RP, Salzemann J, Bloch V, Stockinger H, Berglund AC, Blanchet C, et al. PDB-REDO: Automated re-refinement of X-ray structure models in the PDB. J Appl Crystallogr. 2009;

35. Ehrenmann F, Kaas Q, Lefranc MP. IMGT/3dstructure-DB and IMGT/domaingapalign: A database and a tool for immunoglobulins or antibodies, T cell receptors, MHC, IgSF and MHcSF. Nucleic Acids Res. 2009;

36. Apweiler R, Bairoch A, Wu CH, Barker WC, Boeckmann B, Ferro S, et al. UniProt: The universal protein knowledgebase. Nucleic Acids Res. 2004;

37. Bohne-Lang A, Lang E, Förster T, Von der Lieth CW. LINUCS: LInear Notation for Unique description of Carbohydrate Sequences. Carbohydr Res. 2001;

38. Hamelryck T, Manderick B. PDB file parser and structure class implemented in Python. Bioinformatics. 2003;

39. Cock PJA, Antao T, Chang JT, Chapman BA, Cox CJ, Dalke A, et al. Biopython: Freely available Python tools for computational molecular biology and bioinformatics. Bioinformatics. 2009;

40. Dunbar J, Fuchs A, Shi J, Deane CM. ABangle: Characterising the VH-VL orientation in antibodies. Protein Eng Des Sel. 2013;

41. Giudicelli V, Chaume D, Lefranc MP. IMGT/GENE-DB: A comprehensive database for human and mouse immunoglobulin and T cell receptor genes. Nucleic Acids Res. 2005;

42. Galizia G, Lieto E, De Vita F, Orditura M, Castellano P, Troiani T, et al. Cetuximab, a chimeric human mouse anti-epidermal growth factor receptor monoclonal antibody, in the treatment of human colorectal cancer. Oncogene. 2007.

43. Lee HS, Qi Y, Im W. Effects of N-glycosylation on protein conformation and dynamics: Protein Data Bank analysis and molecular dynamics simulation study. Sci Rep. 2015;

44. Mesters JR, Hilgenfeld R. Protein glycosylation, sweet to crystal growth? In: Crystal Growth and Design. 2007.

45. Davis SJ, Puklavec MJ, Ashford DA, Harlos K, Jones EY, Stuart DI, et al. Expression of soluble recombinant glycoproteins with predefined glycosylation: Application to the crystallization of the t-cell glycoprotein cd2. Protein Eng Des Sel. 1993;

46. Goh JB, Ng SK. Impact of host cell line choice on glycan profile. Critical Reviews in Biotechnology. 2018.

47. Boyd SD, Gaëta BA, Jackson KJ, Fire AZ, Marshall EL, Merker JD, et al. Individual Variation in the Germline Ig Gene Repertoire Inferred from Variable Region Gene Rearrangements. J Immunol. 2010;

48. Liu L, Wang P, Nair MS, Yu J, Rapp M, Wang Q, et al. Potent neutralizing antibodies against multiple epitopes on SARS-CoV-2 spike. Nature. 2020;

49. McCann KJ, Ottensmeier CH, Callard A, Radcliffe CM, Harvey DJ, Dwek RA, et al. Remarkable selective glycosylation of the immunoglobulin variable region in follicular lymphoma. Mol Immunol. 2008;

50. Koers J, Derksen NIL, Ooijevaar-de Heer P, Nota B, van de Bovenkamp FS, Vidarsson G, et al. Biased N-Glycosylation Site Distribution and Acquisition across the Antibody V Region during B Cell Maturation. J Immunol. 2019;

51. Zhu D, McCarthy H, Ottensmeier CH, Johnson P, Hamblin TJ, Stevenson FK. Acquisition of potential N-glycosylation sites in the immunoglobulin variable region by somatic mutation is a distinctive feature of follicular lymphoma. Blood. 2002;

52. Kirkham PM, Schroeder HW. Antibody structure and the evolution of immunoglobulin v gene segments. Semin Immunol. 1994;

53. Shapiro GS, Aviszus K, Ikle D, Wysocki LJ. Predicting regional mutability in antibody V genes based solely on di- and trinucleotide sequence composition. J Immunol. 1999;

54. Wiens GD, Heldwein KA, Stenzel-Poore MP, Rittenberg MB. Somatic Mutation in VH Complementarity-Determining Region 2 and Framework Region 2: Differential Effects on Antigen Binding and Ig Secretion. J Immunol. 1997;

55. Wyatt R, Sodroski J. The HIV-1 envelope glycoproteins: Fusogens, antigens, and immunogens. Science (80-). 1998;

56. Wu Y, West AP, Kim HJ, Thornton ME, Ward AB, Bjorkman PJ. Structural basis for enhanced HIV-1 neutralization by a dimeric immunoglobulin g form of the glycan-recognizing antibody 2G12. Cell Rep. 2013;

57. Calarese DA, Scanlan CN, Zwick MB, Deechongkit S, Mimura Y, Kunert R, et al. Antibody domain exchange is an immunological solution to carbohydrate cluster recognition. Science (80-). 2003;

58. Lee PS, Yoshida R, Ekiert DC, Sakai N, Suzuki Y, Takada A, et al. Heterosubtypic antibody recognition of the influenza virus hemagglutinin receptor binding site enhanced by avidity. Proc Natl Acad Sci U S A. 2012;

59. Pancera M, McLellan JS, Wu X, Zhu J, Changela A, Schmidt SD, et al. Crystal Structure of PG16 and Chimeric Dissection with Somatically Related PG9: Structure-Function Analysis of Two Quaternary-Specific Antibodies That Effectively Neutralize HIV-1. J Virol. 2010;

60. Romijn RAP, Bouma B, Wuyster W, Gros P, Kroon J, Sixma JJ, et al. Identification of the Collagen-binding Site of the von Willebrand Factor A3-domain. J Biol Chem. 2001;

61. Wright A, Tao MH, Kabat EA, Morrison SL. Antibody variable region glycosylation: Position effects on antigen binding and carbohydrate structure. EMBO J. 1991;

